# Decoding the functional role of the calcium ATPase YloB in microbially induced calcite precipitation and sporulation in *Solibacillus silvestris*

**DOI:** 10.1101/2025.09.25.678582

**Authors:** Michael Seidel, Julia Bauer, Carsten Geiß, Susanne Gebhard

**Affiliations:** Institute of Molecular Physiology, Hanns-Dieter-Hüsch-Weg 17, Johannes Gutenberg University, 55128 Mainz, Germany; Institute of Developmental Biology & Neurobiology, Johann-Joachim-Becher-Weg 13, Johannes Gutenberg University, 55128 Mainz, Germany

**Keywords:** endospore formation, heat tolerance, stress resistance, stress adaptation

## Abstract

Calcium homeostasis is essential for bacterial physiology, yet its regulation and role in specialized processes such as microbially induced calcite precipitation (MICP) and sporulation remain poorly understood. Here, we characterized the sole annotated P-type calcium ATPase in *Solibacillus silvestris*, which was previously shown to be upregulated under MICP conditions.

We named the protein YloB based on homology to the *Bacillus subtilis* protein. Structural modeling confirmed that YloB possesses the conserved domains and motifs typical of calcium binding P-type ATPases. Deletion of *yloB* did not affect growth under elevated calcium levels nor did it reduce MICP yields, indicating that YloB is dispensable for calcium detoxification and biomineralization. In contrast, Δ*yloB* cells produced only partially dehydrated spores with severely reduced heat resistance, a defect rescued by complementation. A YloB-mNeonGreen fluorescent fusion localized specifically to the cell membrane surrounding the forespore during early engulfment, and was retained at this site throughout the remainder of sporulation. A transcriptional P_*yloB*_*-msfGFP* reporter demonstrated that *yloB* transcription was induced in the early stages of sporulation, but did not respond to a calcium stimulus.

These results show that YloB does not act to detoxify calcium from the cytoplasm of *S. silvestris*, but rather functions to pump calcium into the forespore, enabling Ca-dipicolinic acid accumulation and proper spore maturation. Our study reveals a specialized role for a bacterial calcium ATPase in sporulation, distinguishing calcium transport for endospore development from biomineralization and global calcium homeostasis.

## Introduction

The calcium concentration in living cells is tightly regulated and plays an important role in signaling (Clapham 2007). While the function of calcium has been extensively studied in eukaryotes, the regulation of calcium homeostasis and its role in bacterial physiology remain to be fully elucidated. Notably, intracellular bacterial calcium levels are kept in a narrow range between 100 to 300 nM, regardless of extracellular conditions, underscoring that calcium homeostasis is carefully controlled (Dominguez 2004). In response to calcium exposure or depletion, the bacterium *Bacillus subtilis* was shown to undergo significant alterations in gene expression, supporting a crucial role of Ca^2+^ ion homeostasis in bacterial physiology (Domniguez 2011). It has also been shown that calcium influences various physiological processes in a range of bacteria, such as swarming motility, virulence and sporulation (Gode-Potratz et al. 2010; Hogarth und Ellar 1978; King et al. 2020).

One process in *B. subtilis* that is well known to require controlled Ca^2+^-levels is sporulation. This developmental cascade is triggered by nutrient depletion and is initiated by phosphorylation of the master regulator Spo0A, which activates genes for asymmetric division of the mother cell to form the forespore. The forespore is subsequently engulfed in a phagocytosis-like process and matures into a highly stress-resistant spore, encased by two membranes, a peptidoglycan layer, and a protective coat (McKenney et al. 2013; Tibocha-Bonilla et al. 2025). As the sporulation process reaches its late stages, the spore becomes increasingly dehydrated through the accumulation of dipicolinic acid (DPA), comprising approximately 10% of dry weight in spores (Paidhungat et al. 2000). DPA is chelated in a 1:1 ratio with mainly calcium ions, ensuring the spore’s stress resistance, especially against wet heat (Church und Halvorson 1959; Magge et al. 2008).

Spore germination is triggered by the sensing of specific signal molecules, including certain amino acids, sugars, and nucleotides that indicate favourable conditions for cell growth. The commitment of the spore to germinate leads to the release of Ca-DPA and the hydrolysis of cortex peptidoglycan (Setlow et al. 2017). This leads to a series of events such as complete spore hydration, an increase in membrane permeability and protein restoration which results in the outgrowth into a vegetative cell (Setlow et al. 2017).

Beyond intracellular processes, calcium was suggested to also promote and stabilize biofilms by the formation of calcite lead structures (Keren-Paz et al. 2022; Nishikawa und Kobayashi 2021). Calcite is a calcium carbonate mineral that can be produced by bacteria in a process called microbially induced calcite precipitation (MICP), which is prevalent throughout the bacterial world (Hoffmann et al. 2021; Boquet et al. 1973). In general, there are two pathways through which MICP can occur, the ureolytic and non-ureolytic pathway. Calcite precipitation by ureolytic bacteria involves the hydrolysis of urea to ammonia and carbon dioxide by the enzyme urease (Stocks-Fischer et al. 1999). This causes an increase in pH and, in the presence of calcium ions, leads to the precipitation of insoluble calcium carbonate in the surrounding medium. On the other hand, non-ureolytic MICP describes a variety of metabolic processes including, for instance, denitrification, sulphate reduction, photosynthesis and heterotrophic pathways, which result in the precipitation of calcium carbonate, although it is not fully understood how these processes lead to mineral formation (Hoffmann et al. 2021; Fahimizadeh et al. 2022). We recently showed that a soil isolate of the species *Solibacillus silvestris* could perform heterotrophic MICP using catabolism of acetate to provide the necessary carbonate ions from the released CO_2_ (Seidel et al. 2025). To provide calcium for MICP, it was proposed that ambient calcium passively enters the cell and is actively exported by calcium pumps. Release of calcium ions from the export system would thereby lead to a local enrichment of the ion on or near the cell surface, creating the supersaturated conditions needed to trigger mineral formation on the bacterial cell surface (Hammes und Verstraete* 2002; Hoffmann et al. 2021; Seidel et al. 2025). For *S. silvestris*, this hypothesis was supported by the observation that the sole P-type calcium ATPase annotated in the genome of the organism was upregulated around six-fold upon calcium supplementation in stationary phase (Seidel et al. 2025).

A different model for MICP has been proposed for *B. subtilis*, where it was reported that, rather than the mineral being formed on or around the cell, calcite formation starts intracellularly by a subpopulation of cells during biofilm formation (Keren-Paz et al. 2022). The authors hypothesized that intracellular accumulation of amorphous calcium carbonate granules, facilitated by P‐type ATPases such as YloB, serves as a nucleation hub. Nucleated minerals are subsequently released to establish an extracellular calcite scaffold, effectively building a rigid skeleton for the biofilm. The study also proposed that biomineralization is an actively regulated mechanism, coupling calcium homeostasis to colony morphogenesis rather than arising from passive precipitation as an indirect consequence of metabolic activity (Keren-Paz et al. 2022). Interestingly, *B. subtilis* YloB shows homology to the P-type ATPase of *S. silvestris* identified as upregulated during MICP-conditions (Seidel 2025).

P-type ATPases are multi-domain, membrane-bound proteins that use ATP to pump ions across biological membranes (Bublitz et al. 2010). They are grouped by the ion they transport; those specific for Ca^2^+ belong to the P2A subclass and actively maintain intracellular calcium homeostasis through ATP‐driven phosphorylation–dephosphorylation cycles. All share a conserved core architecture—three cytoplasmic domains (the phosphorylation domain with a DKTG motif, the nucleotide‐binding domain, and the actuator domain with a TGE motif) plus ten transmembrane helices forming the ion pathway—that undergo conformational shifts to move Ca^2^+ across the membrane (Palmgren und Nissen 2011). To date, only a few calcium P-type ATPases have been shown to be directly implicated in the export of calcium ions from bacterial cells. Two of these calcium ATPases were identified in the human pathogens *Streptococcus pneumoniae* and *Mycobacterium tuberculosis*, where they were found to be essential to cope with toxic calcium concentrations in a eukaryotic host (Rosch et al. 2008; Garg et al. 2020). Beyond their role in exporting calcium from bacterial cells, calcium-binding P-type ATPases have also been found to act as calcium importers during growth in a calcium-deficient environment or for the development of heat-resistant endospores in *B. subtilis* (Raeymaekers et al. 2002; Gupta et al. 2017).

To better understand the calcium flux in *S. silvestris* during the MICP process, we here aimed to characterize the sole calcium P-type ATPase annotated in the genome of the organism, which we named YloB based on its homology to the *B. subtilis* gene. Generation and testing of a *yloB* deletion mutant for growth at increased calcium levels and MICP yields showed no differences to the wild type. However, cells lacking YloB were found to form mostly immature spores that were not fully dehydrated and lacked heat resistance.

Fluorescently labelled YloB clearly localized to the spore membrane during the early steps of spore engulfment, whereas no YloB could be detected in non-sporulating cells. Using a transcriptional reporter assay, we demonstrated that P_*yloB*_ was predominantly active during stationary phase, and activity was specific to cells that had initiated forespore engulfment.

Promoter activation upon the addition of calcium was not conclusively observed. These findings indicate that YloB is vital for correct sporulation, most likely by providing the calcium ions required for spore dehydration, but does not play an active role in calcium detoxification or biomineralization by *S. silvestris*.

## Results

### *In silico* analysis of YloB

To investigate the putative function of the candidate calcium-binding ATPase in *S. silvestris*, encoded in locus AB1K09_00890, we performed a reciprocal BLAST analysis and found that the direct homolog of the protein was YloB in *B. subtilis*, sharing a percentage identity of approximately 43% and a similarity of 64% (Table 1). Therefore, we named the *S. silvestris* protein YloB in accordance with its homolog.

**Table 1.** Reciprocal BLAST reveals *B. subtilis* YloB as the direct homolog of the P-type calcium-transporting ATPase AB1K09_00890 of *S. silvestris*.

| Query | Search genome | Top hit<sup>a</sup> | E-value | Query coverage | % Identity | % Similarity |
| --- | --- | --- | --- | --- | --- | --- |
| *S. silvestris*<br>AB1K09_00890 | *B. subtilis*<br>168 | *yloB* | 0.0 | 96% | 42.81 | 64 |
| *B. subtilis yloB* | *S. silvestris*<br>CGN12 | AB1K09_00890 | 0.0 | 97% | 42.92 | 63 |

<sup>a</sup> In blastp analysis.

A predicted structural model generated with AlphaFold3 (Abramson et al. 2024) revealed that YloB of *S. silvestris* consists of three headpiece domains and ten transmembrane helices and shows the classical five domains of an ATPase: The actuator domain, the nucleotide binding domain responsible for ATP binding, and the phosphorylation domain are all predicted to be cytoplasmic (FIG 1A). The first six transmembrane helices harbour the calcium binding sites and represent the transport domain, while the remaining four helices spanning the membrane form the support domain (FIG 1A) (Kühlbrandt 2004). An overlay with the predicted structure of YloB of *B. subtilis* showed a high degree of similarity, with only minor differences (FIG S1). Next, we conducted a pairwise sequence alignment between the two homologs to check for presence of the conserved residues known to be important for P-type ATPase function (Raeymaekers et al. 2002; Kühlbrandt 2004) (FIG S2). This revealed that the residues involved in ATP binding as well as the phosphorylation site were conserved between the proteins. While the residues for the first calcium binding site are identical between the proteins, two amino acids of the second calcium binding site differ. Specifically, Glu702 and Gln809 in *B. subtilis* are substituted with Ala680 and Arg785 in *S. silvestris*, respectively, which likely results in a net loss of negative charge at the first calcium binding site (FIG S2), the consequences of which are unclear. To conclude, these findings indicate that ΔYloB of *S. silvestris* likely is a functional calcium-binding ATPase.

**Figure 1.**
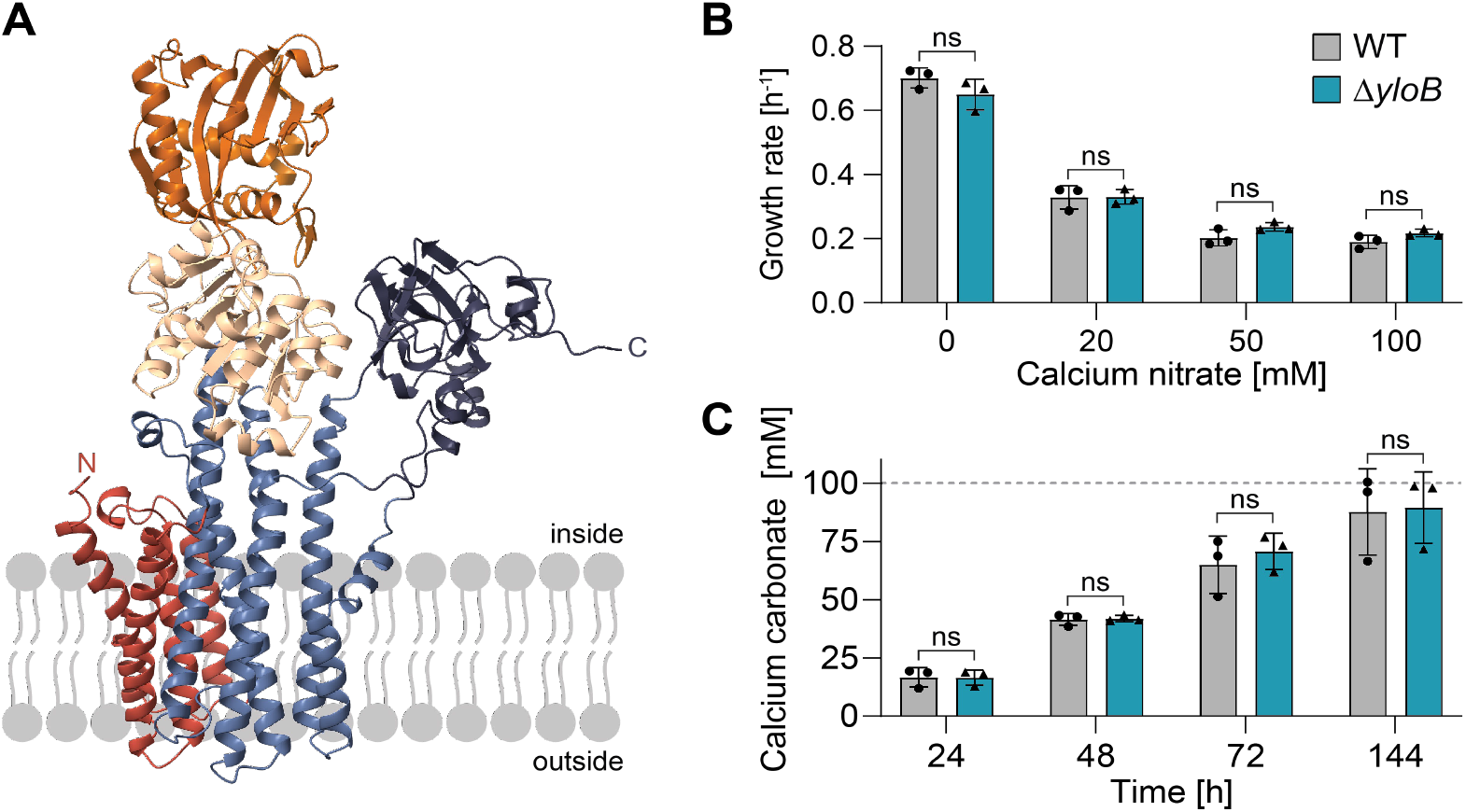
YloB is a calcium-binding ATPase, but dispensable for growth and calcium carbonate precipitation. (A) Structure prediction was generated by AlphaFold3 (Abramson et al. 2024) and visualized using ChimeraX (Meng et al. 2023). The protein’s N- and C-termini are labelled. The orientation of the protein was determined by 2D topology using PROTTER analysis (Omasits et al. 2014). The three cytoplasmic headpiece domains were identified as the actuator domain (dark blue), the nucleotide binding domain (orange) and the phorylation domain (beige), while the transport (light blue) and support (red) domains span the membrane (Kühlbrandt 2004). (B) Growth rate during exponential phase of indicated strains grown at different calcium nitrate concentrations. (C) Precipitated calcium carbonate of indicated strains. All cultures were grown in YAC medium for the indicated times, when the calcium carbonate precipitate was harvested and quantified. The horizonal dashed line visualizes the theoretical maximum of calcium carbonate precipitation based on 100 mM available calcium nitrate. For panels B and C, the bars depict the mean, error bars are the standard deviation of the mean, and individual data points of independent triplicate experiments are shown as black symbols. No significant (ns) differences between the WT and Δ*yloB* strains were observed in a Student’s t-test.

### Deletion of *yloB* does not affect calcium sensitivity of *S. silvestris*

To investigate the physiological role of YloB, we deleted the gene by double homologous recombination using a temperature-sensitive pMAD-oriT-derived plasmid (Seidel et al. 2025). We readily obtained the deletion mutant (Δ*yloB*), suggesting that the gene is not essential.

To determine if YloB is involved in calcium homeostasis in *S. silvestris*, we determined the growth rate of the wild type (WT) and Δ*yloB* mutant upon calcium stress, supplementing the growth medium with 0, 20, 50, and 100 mM calcium nitrate. Without any calcium stress, WT cells showed a growth rate of 0.70 h^-1^. We observed that already the addition of 20 mM calcium nitrate led to a significant decrease of its growth rate to 0.33 h^-1^, with a further reduction to 0.20 h^-1^ when cultivated with 50 mM calcium nitrate (FIG 1B, S3). However, increasing the calcium nitrate concentration from 50 mM to 100 mM did not result in a further decline. In all tested conditions, Δ*yloB* showed no significant difference to WT growth behaviour (FIG 1B, S3).

### YloB is dispensable for biomineralization

In previous studies, it was hypothesized that YloB might be involved in the MICP process in *B. subtilis*, and it was shown that *yloB* expression was upregulated in MICP-conditions in *S. silvestris* (Keren-Paz et al. 2022; Seidel et al. 2025). To determine whether YloB plays an active role in MICP in *S. silvestris*, WT and Δ*yloB* cells were cultivated in YAC precipitation medium (100 mM calcium nitrate). Quantification of mineralised calcium carbonate after one to six days showed that the amount of mineral formed increased steadily over time (FIG 1C). After six days, 90% of the available soluble calcium had precipitated as mineral in both WT and Δ*yloB* cultures, with no statistically significant differences observed between strains (FIG 1C). This indicated that YloB was not required for biomineralization by *S. silvestris* and therefore likely is not responsible for actively providing Ca-ions for mineral nucleation.

### The calcium-transporting ATPase YloB affects sporulation in *S. silvestris*

As YloB in *B. subtilis* was shown to be important for the dehydration of endospores (Raeymaekers et al. 2002), we analysed sporulation of *S. silvestris* WT and Δ*yloB* cells. For this, we imaged the cells by phase contrast microscopy after 24 h and 48 h of growth in YA medium. In total, around 3% of both WT and Δ*yloB* cells showed a developing spore after 24 h (Fig. S4), which increased to approximately 5% after 48 h with no difference observed between the two strains (FIG 2A). A striking difference was observed in the spore morphology of the two strains, with the WT forming larger spores with stronger phase-brightness in comparison to Δ*yloB* (FIG 2B), suggesting that Δ*yloB* spores might not fully mature. To test whether this morphological difference resulted in a defect of heat-resistance, we determined the proportion of viable cells (CFU/ml) remaining after pasteurization of the cultures. While 0.89% of all WT cells had formed a heat-resistant spore after 48 h, Δ*yloB* showed a 3-log reduction, with only 0.0009% of all cells having formed a heat-resistant spore (FIG 2C).

**Figure 2.**
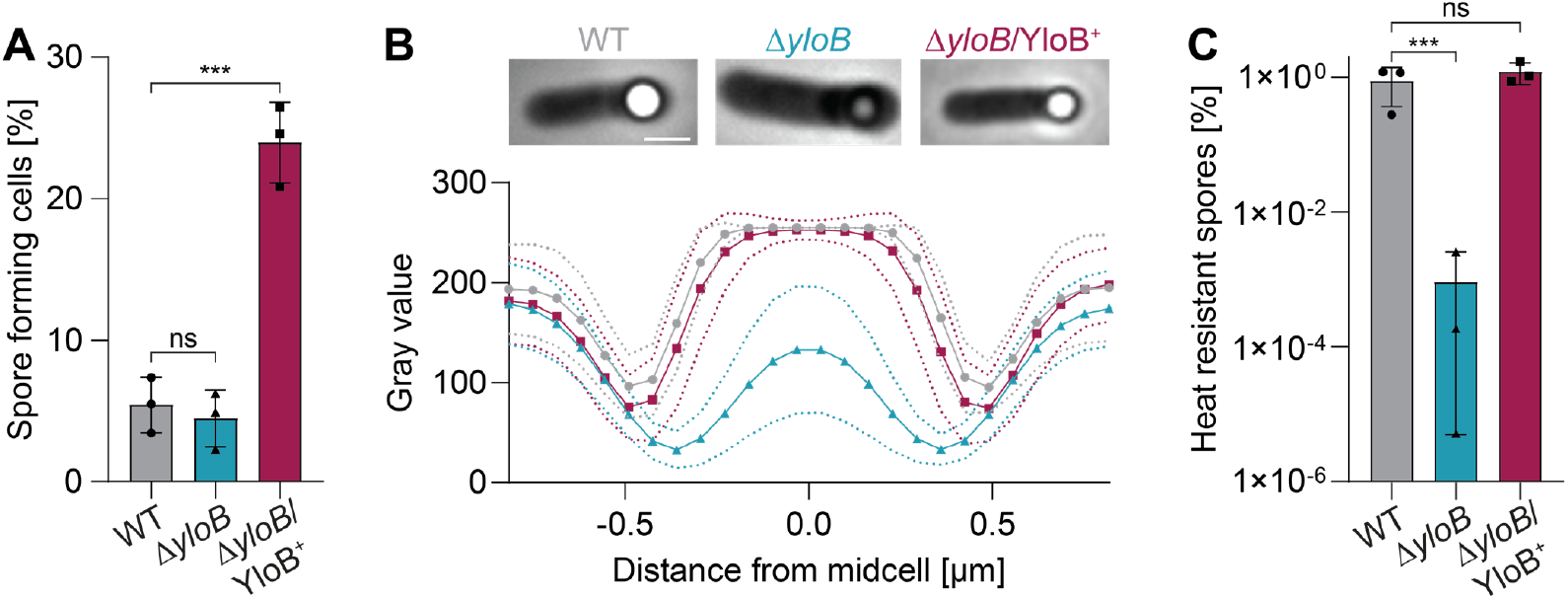
Cells lacking YloB show a severe sporulation defect. (A) Proportion of total spores of indicated strains determined by phase contrast microscopy after 48h of growth in YA medium. The fraction of sporulated cells was divided by the total cell count. Student’s t-test was conducted to test for significant differences. The mean and the standard deviation for biological triplicates are represented. ns=not significant, p > 0.005 (***). (B) Top: Representative phase contrast images of sporulated cells of the indicated strains. Bottom: Spore brightness was quantified by plotting gray value intensity profiles across the width of the cell, n ≥31. Symbols indicate mean gray values, and dotted lines show standard deviation for each strain. (C) Heat resistance of the spores. The ratio of heat-resistant spores was determined by dividing the viable count (CFU/ml) of the pasteurized culture by that of an untreated sample of the same culture. In panels A and C, the mean (bar), the standard deviation (error bar) and individual data points (symbols) are shown for biological triplicates. Student’s t-test was conducted to test for significant differences. ns=not significant, p > 0.005 (***).

To ensure the difference in spore morphology and loss of heat resistance were indeed due to the absence of YloB, the Δ*yloB* strain was complemented by plasmid-borne expression of an intact copy of *yloB* under control of its native promoter. The resulting strain *yloB*/YloB^+^ showed a spore morphology and proportion of heat-resistant spores that were indistinguishable from the WT (Fig. 2B&C). Curiously, the fraction of sporulated cells in the population was significantly increased in the complemented strain (Fig. 2A), likely caused by the additional copies of the gene on the plasmid construct leading to overproduction of YloB beyond physiological levels. It is not clear why this might lead to more cells being detected as sporulated, but as the proportion of heat-resistant spores was not altered compared to the WT strain, presence of multiple *yloB* copies did not appear to significantly affect final sporulation outcome of the population. Taken together, the complementation experiments showed that the sporulation defects observed in the deletion strain were specifically due to loss of *yloB*, strongly supporting a role of the Ca-ATPase in spore maturation of *S. silvestris*.

### YloB is localized in the spore membrane of *S. silvestris*

Next, we aimed to study the localization of YloB in vegetative cells and during sporulation. To this end, we generated an mNeonGreen (mNG) translational fusion to the C-terminus of YloB. The strain was generated by integrating the fusion into the native gene to circumvent any artefacts from altered gene copy number or changes in gene expression. Microscopic analyses showed that the strain harbouring the mNG-fused YloB protein was able to produce fully dehydrated spores (Fig. 3), and thus the YloB-mNG protein was deemed functional.

**Figure 3.**
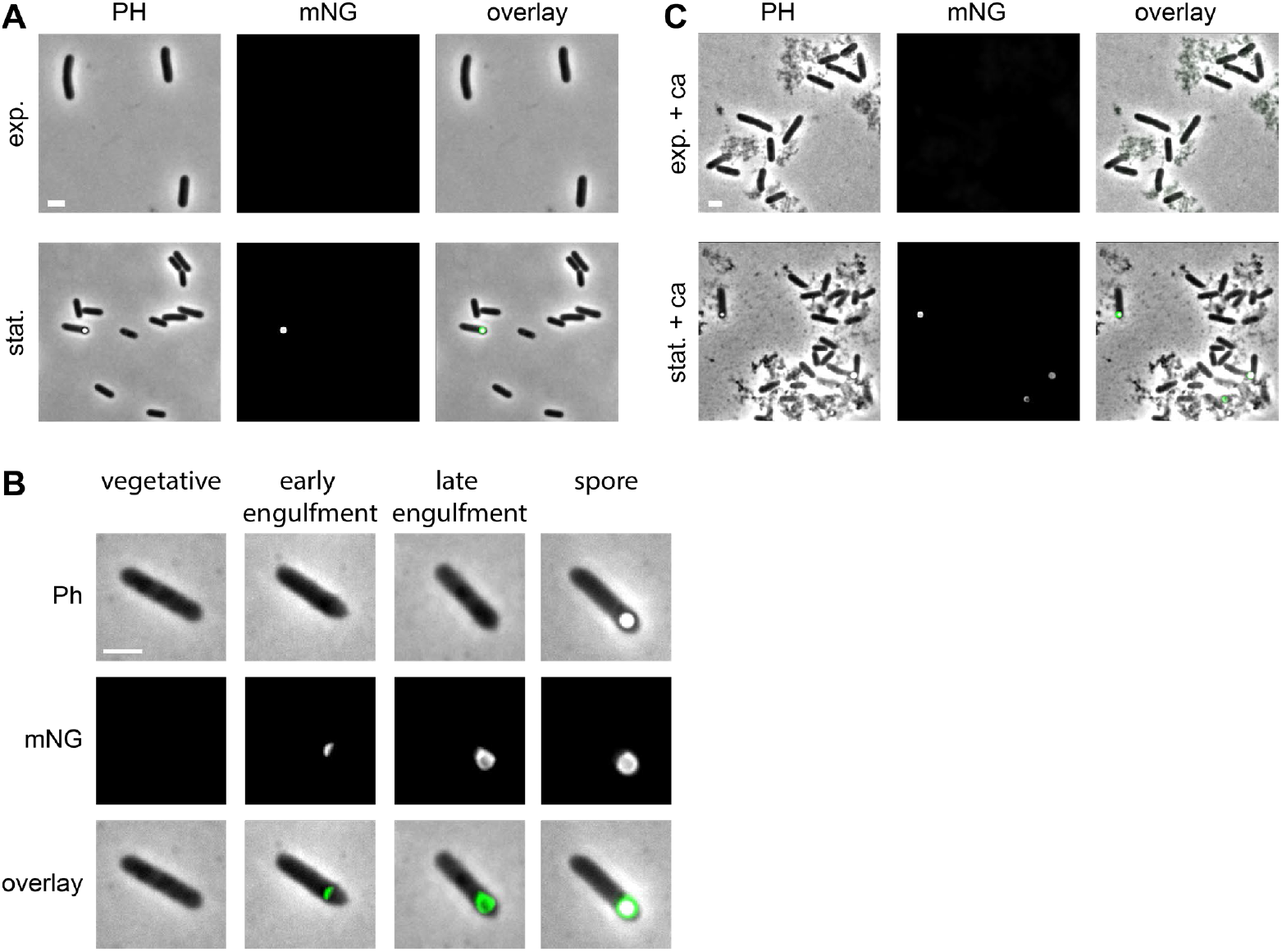
YloB exclusively localizes to the spore membrane. Fluorescence microscopic analyses of *yloB::yloB-mNeonGreen* cells, encoded in the native site. Phase contrast (Ph), mNeonGreen fluorescence (mNG) and the overlay are depicted. Scale bars 2 µm. (A) Cells imaged in exponential and stationary phase in YA medium. (B) Localization of YloB in the spore membrane at different stages of sporulation including vegetative, early and late stage of engulfment of the forespore and dehydrated spore. Sporulation phases were assigned based on phase-contrast morphology. (C) Cells exposed to 100 mM calcium nitrate for 1 h during exponential or stationary phase.Addition of calcium nitrate caused a precipitate, visible as grey specks in the background of the cells.

The mNG fusion strain next was examined by fluorescence microscopy in both exponential and stationary growth phases. No fluorescent signal was detected in exponentially growing cells. For cells in stationary phase, most cells did not show a fluorescent signal, but in cells containing a spore, a clear fluorescent signal was seen surrounding the spore (FIG 3 A&B, bottom panel), suggesting that YloB was only produced in sporulating cells and located to the membrane environment around the spore. Consistently with this, fluorescence was also observed in cells in the earlier phases of sporulation, i.e. engulfment of the forespore. These cells showed a bulge at one cell pole in phase contrast microscopy, without a mature, phase bright spore being visible. In these cells, the YloB-mNG fusion was clearly localized to the region where the mother cell membrane migrates around the forespore, with some cells showing a small fluorescent crescent at the division septum (likely early phase of engulfment), while others showed fluorescence around the whole forespore (late engulfment) (Fig. 3B). Interestingly, no fluorescent signal was observed in the cell membrane away from the division septum, suggesting that YloB possesses an as yet unidentified mechanism to specifically localize to the membrane region engulfing the forespore.

We had previously shown that exposure of stationary phase cells of *S. silvestris* to 100 mM calcium nitrate resulted in a six-fold increase in gene expression of *yloB* determined by RT-qPCR (Seidel et al. 2025). To further explore this, we supplemented both exponential and stationary phase cells with 100 mM calcium nitrate for 1 h and examined whether this treatment altered the overall fluorescence intensity or localization of YloB-mNG. However, no clear differences were observed compared to untreated controls (Fig. 3A&C).

### P_*yloB*_ is active during sporulation

Given our observation that the mNG-tagged protein was only produced in sporulating cells, we next wanted to test whether the previously observed change in gene expression was indeed specifically due to calcium addition or an unspecific effect caused by the chosen stress condition. To test this, we constructed a transcriptional reporter strain, where the native promoter of *yloB* (P_*yloB*_) was placed in front of the gene encoding monomeric superfolder GFP (msfGFP). The resulting plasmid construct (pMAD-oriT-P_*yloB*_*-*msfGFP) was introduced into WT *S. silvestris* by conjugation. Because the standard growth temperature of this bacterium is within the permissive range for replication of the vector, this allowed stable, extrachromosomal maintenance of the construct.

Flow cytometry analysis revealed that the promoter was highly active in stationary phase cells, with about 82% of the bacterial cells exhibiting a GFP signal (FIG 4A, Fig. S5). In contrast, only approximately 5% of the cell population in the exponential phase were detected as GFP positive. This indicated that the *yloB* promoter was predominantly active during stationary phase, and almost completely inactive during exponential phase, consistent with our observations for production of the YloB-mNG protein fusion shown above.

**Figure 4.**
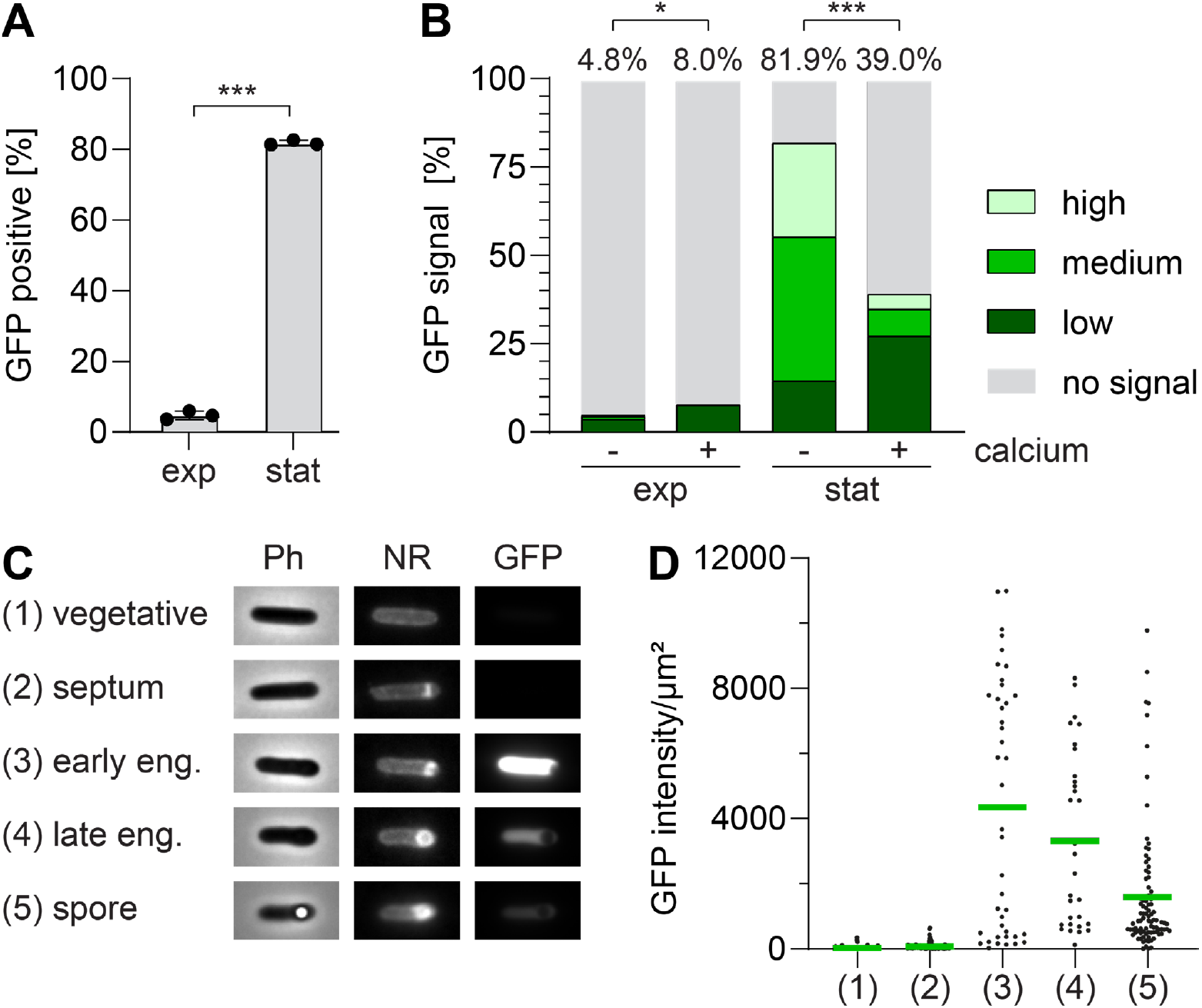
*PyloB* shows growth phase-dependent activation and is repressed by a high calcium level in stationary phase. (A) Cells were cultivated to the exponential (exp) and stationary (stat) phase and the proportion of GFP positive cells was determined using flow cytometry, with a negative control strain harbouring promoter-less msfGFP used to set the cut-offs. The mean and the standard deviation for biological triplicates are represented. (B)Quantification of GFP signal intensities. Cells were cultivated to the exponential or stationary growth phase and challenged with or without 100 mM calcium nitrate for 1 h, as indicated. The proportion of GFP positive cells was quantified by flow cytometry, with GFP-positive cells further divided into those with high, medium or low signal intensity, shown by coloured bar segments. The total proportion of GFP positive cells is shown in % on top. Data represent the mean of three biological replicates. (C) Microscopic visualization of the GFP-reporter strain in stationary phase at the indicated stages of sporulation. Before microscopy, cell membranes were stained with NileRed(NR) for 5 min. Sporulation stages were assigned based on cell morphology in phase contrast (Ph) images and with NR staining. GFP fluorescence was then assessed for each stage. (D) Quantification of GFP intensity per μm^2^ over different stages of sporulation as shown in C. Student’s t-test was conducted to test for significant differences, p > 0.01 (*), p > 0.005 (***).

Moreover, the flow cytometric data showed a distinct grouping of the GFP-positive cells into three sub-populations, which we used to group the cells into three categories of low, medium or high fluorescence intensity (Fig. 4B). This highlighted further differences between the growth phases. During exponential phase, the low proportion of cells for which a GFP signal was detectable predominantly fell into the “low intensity” category, supporting the lack of promoter activity during exponential growth. Cells with medium or high fluorescence intensity were almost exclusively detected during stationary phase and together accounted for the majority of GFP-positive cells in this growth phase, again consistent with the results obtained for the YloB-mNG protein fusion.

With functionality of the transcriptional reporter established, we next wanted to see whether exposure to calcium elicited changes in P_*yloB*_ activity. Following challenge with 100 mM calcium nitrate for 1 h, exponential phase cells showed a small but statistically significant increase from around 5% to 8% in GFP positive cells, with most of these cells still falling into the low fluorescence category (FIG 4B). For stationary phase cells, we observed a marked decrease in overall GFP-positive cells from around 82% in the control to around 39% after calcium challenge. In addition, the distribution of fluorescence intensity shifted from predominantly medium or high in the untreated culture to a mainly low intensity signal for the culture supplemented with calcium (FIG 4B). This was surprising given we had previously observed elevated *yloB* mRNA levels during MICP-conditions (Seidel et al. 2025). One explanation might be that the calcium shock caused a reduction in cells initiating sporulation, thus requiring less P_*yloB*_ activity. However, indirect effects due of the increased calcium level, e.g. slowed msfGFP protein folding, might also have caused a reduced fluorescence output, thus definitive conclusions cannot be drawn at this stage. Nevertheless, the data clearly suggest that P_*yloB*_is not specifically activated by the presence of high calcium concentrations. This in turn indicates that the observed change in *yloB* mRNA detected by RT-qPCR in our earlier study (Seidel et al. 2025) was likely an indirect consequence of the chosen stress conditions.

To understand the basis for the different categories of fluorescence intensities among the GFP-positive cells, we returned to fluorescence microscopy to analyse the P_*yloB*_ reporter strain on the single cell level. To allow us to correlate promoter activity with the different sporulation stages, the cells were stained with the membrane dye Nile Red (NR). Based on the NR staining pattern, cells were categorized into five morphological stages: (1) vegetative cells (even NR staining outlining the cell); (2) septum formation (an asymmetric division septum is visible); (3) early engulfment (a crescent of NR-stained membrane partially surrounds the forespore); (4) late engulfment (a bright NR signal surrounds the entire forespore); and (5) cells containing a dehydrated spore (the NR signal surrounds a phase-bright spore) (FIG 4C). Based on these criteria, each cell was assigned to one category, followed by quantitative analysis of GFP fluorescence per µm2 of the individual cells (Fig.4D). This analysis revealed no or minimal GFP intensity in vegetative cells and those displaying an asymmetric septum. A striking increase in GFP intensity was observed in cells in the early engulfment phase, suggesting that P_*yloB*_ activity is strongly induced at this stage of the sporulation process. Cells in the later sporulation phases, i.e. undergoing late engulfment or containing a dehydrated spore exhibited a reduced, although still clearly visible fluorescence intensity (FIG 4CD). These results indicate that *yloB* expression is specifically induced in the mother cell upon initiation of forespore engulfment, and likely switched off again soon after, explaining the gradually decreasing GFP intensities in cells of the later sporulation stages. Moreover, comparing the microscopic data to the flow cytometric analyses suggests that the three sub-populations with different GFP intensities shown in Fig. 4B likely reflect cells in the different sporulation phases as shown in panels C and D, with the high fluorescence sub-population likely including cells in the forespore engulfment phase, while cells in later sporulation phases or only just beginning forespore engulfment would likely fall into the medium or low intensity category. Taken together, the reporter analyses as well as the YloB-mNG protein fusion show a dynamic regulation *yloB* expression that is tightly linked to sporulation in stationary phase cells, consistent with the marked sporulation defect of the *yloB*-deleted strain.

## Discussion

Calcium is a universal signaling ion that plays vital roles not only in eukaryotes but also in bacteria. Hence, cells need to tightly regulate their calcium homeostasis, but the detailed mechanism by which bacteria achieve this is not understood. In addition, active calcium transport was proposed to be an important factor for biomineralization of calcium carbonate, and it is known to be essential for production of heat-resistant endospores. *S. silvestris* CGN12 is an environmental isolate from a high-calcium natural site (limestone rock), capable of growth at high calcium concentrations, sporulation and calcite biomineralization (Reeksting et al. 2020; Seidel et al. 2025). It therefore presented an ideal model to gain deeper insights into the role of calcium transport in these processes. Here, we set out to characterize the sole annotated calcium P-type ATPase in this bacterium.

*In silico* analyses showed that the calcium ATPase of *S. silvestris* is a homolog of YloB in *B. subtilis*. While sequence identity (43%) and similarity (63%) were only moderate, structural predictions showed a very high similarity between the two proteins (Table 1, FIG S1). This is consistent with the fact that P-type ATPases show high variability in the amino acid sequence, while the overall structure is conserved (Kühlbrandt 2004). All of the key motifs for calcium P-type ATPases were identified in YloB of *S. silvestris*, with the exception of two differing residues in the second calcium binding motif differ, with Glu702 and Gln809 of the *B. subtilis* protein substituted with Ala680 and Arg785 in *S. silvestris*. A Glu-to-Arg exchange was also observed in the P-type ATPase LMCA1 in *Listeria monocytogenes*, where it was found to be responsible for a higher pH optimum of the protein (Faxén et al. 2011). Given that *S. silvestris* optimally grows at alkaline pH (>pH 8), it is plausible that YloB carries similar substitutions to ensure a higher pH optimum. From these results, we concluded that YloB shows all features of a functional calcium ATPase.

To determine the physiological role of YloB in *S. silvestris*, we deleted the encoding gene by double homologous recombination. Since it was hypothesized that YloB could act as a calcium export system in *S. silvestris*, it was expected that its gene deletion would result in a higher sensitivity to calcium compared to the wild type. However, Δ*yloB* cells exhibited comparable sensitivity to calcium as the wild type (FIG 1B). Similar results have been reported for the *yloB* deletion mutant in *B. subtilis* (Keren-Paz et al. 2022), indicating that YloB does not function as a calcium efflux detoxification mechanism. Next, we wondered whether cells lacking YloB could still precipitate calcium into calcite mineral. This hypothesis was supported by previous findings, where increased *yloB* mRNA levels were found under MICP conditions (Seidel et al. 2025). In *B. subtilis*, the deletion of *yloB* was reported to abolish calcite mineral formation during biofilm formation, and thus cause a defect in biofilm formation and architecture (Keren-Paz et al. 2022). However, in *S. silvestris*, the Δ*yloB* strain exhibited a similar calcite precipitation rate and yield as the WT (FIG 1C), indicating that it plays no role in biomineralization. Moreover, we did not observe increased activity of P_*yloB*_ in the transcriptional reporter assays, nor an increased production or altered localization of YloB-mNG in response to increased calcium supplementation. From these data we conclude that YloB is not the transporter responsible for calcium efflux in *S. silvestris*.

Instead, the role of YloB may be more specific to calcium handling during sporulation, where precise calcium delivery is critical for spore maturation. Calcium is required as a chelator for DPA, and the accumulation of Ca-DPA causes partial dehydration of the endospore, which gives the spore heat-resistant properties (Paidhungat et al. 2000). *S. silvestris* cells that lacked YloB produced only partially phase-bright spores, suggesting that they are not fully dehydrated (FIG 2B). While WT and Δ*yloB* cells exhibited similar sporulation rates in terms of total number of sporulating cells, the proportion of generated heat-resistant spores was dramatically reduced in Δ*yloB* (FIG 2A&C). In *B. subtilis* Δ*yloB*, phase-dark spores where shown to have a deficiency in calcium accumulation into the spore core, as calcium levels in the forespore were decreased compared to WT spores (Chen et al. 2019). It is therefore plausible that YloB in *S. silvestris* similarly functions to transport calcium into the forespore.

By studying the activity of the *yloB* promoter as well as the localization of fluorescently tagged YloB-mNG, we could show that YloB has a major role during sporulation. While P_*yloB*_ was active in a small minority of vegetative cells, no YloB-mNG signal could be observed here (FIG 3A, 4A). This is in agreement with observations in *B. subtilis*, where YloB could only be found in sporulating but not in vegetative cells (Raeymaekers et al. 2002). Single-cell measurements by microscopy showed that *S. silvestris* cells that have started forming an asymmetric spore septum strongly activated P_*yloB*_, with declining promoter activities in the later stages of sporulation (FIG 4C&D). At the same early stage of sporulation, the YloB-mNG protein appears specifically at the asymmetric division septum and then follows the forespore engulfment process (FIG 3B). Interestingly, despite the high promoter activity and strong fluorescent signal from the mNG-tagged protein, we could not observe the protein elsewhere in the mother cell membrane (FIG 3A&B). Such a pattern of protein incorporation has been reported for some early-stage spore coat proteins and for SpoVV, a DPA transporter, which are expressed in the mother cell and located in the outer spore membrane (McKenney and Eichenberger 2012; Ramírez-Guadiana et al. 2017). The P_*yloB*_-msfGFP reporter gave a clear signal only in the mother cell cytoplasm, not the spore (FIG 4C), suggesting promoter induction is mother cell-specific and that therefore YloB is produced by the mother cell. How the protein specifically localizes to the division septum and the membrane region surrounding the forespore is an interesting question. One potential explanation could be curvature dependent localization. The membrane growing around the forespore is the only region with positive membrane curvature in the cell, and this has been proposed to target sporulation-relevant proteins to their specific site (Ramamurthi and Losick 2009). An alternative mechanism could be localization through interaction with a partner protein in the inner spore membrane. Such a mechanism has been reported for *B. subtilis* and *Clostridioides difficile* SpoIIIAH, which is located in the outer forespore membrane, but specifically interacts with SpoIIQ in the inner forespore membrane to form a channel through both membranes, facilitating communication between the two cells (Meisner et al. 2008; Serrano et al. 2016; Crawshaw et al. 2014). While further research will be required to elucidate the localization mechanism for YloB, the latter model might be attractive, given that the Ca^2+^-ions will have to cross both the outer and inner forespore membrane to reach the forespore, while YloB can only span one lipid bilayer.

Taken together, our findings suggest that YloB plays a central role in transporting calcium from the mother cell into the forespore to enable proper Ca-DPA accumulation and the formation of heat-resistant spores, rather than serving as a calcium efflux pump for detoxification or involvement in MICP.

## Material & Methods

### Strains and growth conditions

All strains and plasmids used in this study are listed in Table S1. For routine growth, *E. coli* strains were grown in standard lysogeny broth (LB) at 37°C with shaking (120 rpm). Selection for plasmid uptake and maintenance in *E. coli* was achieved with 100 µg/ml ampicillin. Cultures of the conjugation donor strain were additionally supplemented with 50 µg/ml of 5-aminolevulinic acid (5-ALA).

*S. silvestris* was routinely grown in LB pH 8.2, adjusted with 20 mM Tris-HCl pH 9, at 30°C with shaking (120 rpm). To select for successfully conjugated clones carrying the plasmid, LB pH 8.2 medium was supplemented with MLS antibiotics (0.5 µg/mL erythromycin and 12.5 µg/mL lincomycin). Where indicated, *S. silvestris* was cultivated in Yeast extract Acetate (YA) medium containing 50 mM Tris-HCl pH 7.8, 0.2% w/v yeast extract, 0.5 mM MgSO_4_, 0.01 mM MnSO_4_ and 200 mM sodium acetate. The precipitation medium YAC was based on YA medium and additionally contained 100 mM calcium nitrate, unless otherwise stated.

### Microbially induced calcite precipitation (MICP) assay

Overnight cultures grown in LB medium were used to inoculate 20 ml YAC media to an initial OD_600_ of 0.02. The cultures were incubated for 1 to 6 days at 30°C and shaking at 140 rpm.

The cultures were then harvested by centrifugation at 400 x *g* for 5 min. To remove bacterial cells from the precipitate the pellet was repeatedly washed with distilled water until the supernatant remained clear. The precipitate was dried at 60°C until no further change in weight was detectable. The amount of precipitate was determined by calculating the difference in weight of the centrifuge tubes before the experiment and after drying the precipitate. The concentration of precipitated calcium carbonate in the original culture was calculated from the weight of the precipitate, molecular mass of CaCO_3_ and the harvested culture volume.

### Plasmid construction

Genomic DNA of *S. silvestris* was used to amplify genomic regions of interest. All generated plasmids are derivates of pMAD-oriT (Seidel et al. 2025) and were constructed via Gibson Assembly (Gibson et al. 2009) with minor modifications. A 5x isothermal reaction buffer was prepared containing 500 mM Tris-HCl (pH 7.5), 50 mM MgCl_2_, 1 mM dNTPs, 50 mM DTT, 25% (w/v) PEG-8000, and 5 mM NAD. The Gibson Assembly Master Mix (1.2 mL total) was set up with 1x isothermal reaction buffer, 6.4 units T5 exonuclease, 40 units Phusion DNA polymerase, and 6.4 units Taq DNA ligase, and aliquoted into 15 µL portions. Assembly reactions were set up by combining a master mix aliquot and 300 ng linearized vector with a 1:3 molar ratio of insert DNA in a final volume of 20 µL, followed by incubation at 50°C for 30 min. Subsequently, 10 µL of the reaction was used to transform competent *E. coli*.

Linearised vector was generated either by restriction digestion via the EcoRI/SalI sites or by PCR using the primer pair SG1680 and SG1681, followed by a DpnI digest to remove the template DNA. Correct assembly was verified by colony PCR using the primer pair SG1540 and SG1541, followed by sequencing of the insert.

#### pMSMAD03

Plasmid to delete *yloB* via double homologous recombination. To amplify the upstream fragment, the primers SG1682 and SG1684 were used. To generate the downstream fragment, the primers SG1683 and SG1685 were used. The final insert fragment was obtained via overlap PCR using the primers SG1686 and SG1687 and then used for Gibson Assembly with linear pMAD-oriT.

#### pMSMAD05

Plasmid for insertion of *mNeonGreen* as a C-terminal fusion to *yloB* via double homologous recombination. To amplify the upstream fragment, the primers SG1694 and SG1695 were used. To obtain the *mNeonGreen* fragment, the primers SG1696 and SG1697 were used using pSHP1 as the template. To generate the downstream fragment, the primers SG1698 and SG1699 were used. The final insert fragment was obtained via overlap PCR using the primers SG1694 and SG1699 and then used for Gibson Assembly with linear pMAD-oriT.

#### pMSMAD06

Plasmid used for the reporter assay as promoter-less *sfGFP* control. To amplify *sfGFP*, the primer pair SG1796 and SG1787 were used, using pHJS105 as a template. The resulting fragment was then used for Gibson Assembly with linear pMAD-oriT.

#### pMSMAD08

Plasmid used for the reporter assay, containing P_*yloB*_ *sfGFP*. To amplify the promoter of *yloB*, the primers SG1790 and SG1795 were used. The resulting fragment was used in a Gibson Assembly reaction together with pMSMAD06 that was linearized using the primers SG1793 and SG1681.

#### pMSMAD11

Plasmid for complementation of *yloB* under its native promoter. The fragment P_*yloB*_ *yloB* was obtained using the primers SG1708 and SG1820 and used for Gibson Assembly with linear pMAD-oriT.

### Construction of in-frame DNA deletions and insertions in *S. silvestris*

For the incorporation of constructed vectors into *S. silvestris*, conjugation was performed as described (Seidel et al. 2025), with the exception of using *E. coli* ST18 as the donor strain to facilitate counterselection based on this strain’s auxotrophy for 5-ALA (Thoma and Schobert 2009). Overnight cultures of *S. silvestris* and transformed *E. coli* ST18 were prepared by inoculating 5 ml LB medium at the respective pH values stated above, with addition of 50 µg/ml 5-ALA for ST18. The following day, for each strain 20 ml of LB medium plus 10 mM of MgCl_2_, (and with 5-ALA for ST18) were inoculated to an initial OD_600_ of 0.1, and incubated until ST18 reached the exponential phase (OD_600_ = 0.4-0.6). The bacterial cells were pelleted by centrifugation, and ST18 cells were additionally washed in 1 ml LB to remove residual 5-ALA. Both cell pellets were resuspended and combined in a total volume of 10 ml standard LB medium containing 20 mM MgCl_2_. After incubation for 1 h at 30°C, the cells were again pelleted and resuspended in the small amount of medium remaining after decanting the supernatant. The cell suspension was spotted on an LB plate containing 20 mM MgCl_2_ and incubated for 24 h at 30°C. The next day, the cells were scraped off and resuspended in 1 ml LB pH 8.2, serially diluted to 10^-2^, and 3 x 20 µl per dilution were spotted on an LB plate containing MLS selection, and incubated at 30°C for up to 2 days until the appearance of colonies.

Deletion of the *yloB* gene and integration of the *yloB*-mNG fusion construct were generated by double homologous recombination as described (Seidel et al. 2025). Briefly, an overnight culture was prepared in LB medium with MLS selection and grown at 30°C. The next day, 10 mL LB with MLS selection was inoculated to a starting OD_600_ of 0.1 and incubated at 30°C for 2 h, at which time the temperature was increased to 42°C for 5 h. Serial dilutions were then plated onto LB agar with MLS selection and incubated at 42°C for 24 h. Individual colonies were selected and used to inoculate 5 ml LB pH 8.2 without antibiotics and incubated overnight at 30°C. The temperature was then increased to 42°C for 3 h. Serial dilutions were plated onto LB pH 8.2 agar without selection and incubated for 24 h at 42°C. Colonies were replica‐patched onto MLS and antibiotic-free plates. MLS sensitive clones were screened for gene deletion or *mNG* insertion by PCR using primers SG1842 and SG1843.

### Growth rate analysis

The growth rate was determined in YA medium with increasing concentrations of calcium nitrate (0 mM, 20 mM, 50 mM and 100 mM). Test tubes of 16 mm diameter containing 3 ml medium were inoculated with an overnight culture to an initial OD_600_ of 0.05 and incubated at 30°C with shaking (120 rpm). The optical density was measured in 30-minute intervals for 7 h using an Ultrospec 10 cell density meter (Biochrom). The exponential growth phase was identified and the growth rate calculated by fitting the data with the exponential growth equation using GraphPad Prism 10.

### Quantification of sporulation and heat-resistance

To determine the heat-resistance of spores in *S. silvestris* WT and the Δ*yloB* strain, 5 ml YA were inoculated with an overnight culture to a starting OD_600_ of 0.05. The cultures were incubated at 30°C for 24 h and 48 h with shaking (120 rpm). The cultures were analyzed by phase contrast microscopy using a Leica DMi8 inverted microscope and a Leica K8 Scientific CMOS camera. For each of three biological replicates, cells from around 7 independent microscopic fields of view were counted (total across all replicates n ∼ 1000). Cells were scored as sporulated if they showed a spore that was at least partially phase-bright. The fraction of sporulated cells was determined by dividing the number of sporulated bacteria by that of the total cell count.

To determine the proportion of heat-resistant spores, serial dilutions were prepared and then pasteurized at 80°C for 20 min. 100 µl of pasteurized and non-pasteurized dilutions were each plated onto LB pH 8.2 plates and incubated overnight at 30°C. Colony numbers were enumerated for the dilution giving between 50 and 300 colonies, and the CFU/ml was determined for both the pasteurized and non-pasteurized cultures. The proportion of heat-resistant spores was determined by dividing the CFU/ml of the pasteurized culture by that of the non-pasteurized culture.

### Fluorescence microscopy and membrane staining

Microscopy was performed with cells immobilised on Teflon-coated multi-spot microscope slides (Hendley-Essex) with 1.5% (w/v) agarose in H_2_O. Per well, 0.5 µl of the respective cell culture was spotted and covered with a cover slip. If required, membranes were stained with NileRed (9-diethylamino-5-benzo[α]phenoxazinone). The staining was conducted by incubation of 100 µl culture with 1 µg/ml of NileRed for 5 min at 30°C, immediately followed by spotting the cells on the agarose covered slide and microscopy.

For imaging cells in exponential phase, 3 ml YA medium were inoculated from an overnight culture to a starting OD_600_ of 0.05, grown to an OD_600_ of 0.5 and then spotted for microscopy, or supplemented with 100 mM calcium nitrate for 1 h and then spotted. For cells in stationary phase, 5 ml YA medium were inoculated from an overnight culture to a starting OD_600_ of 0.05 and grown for either 24 h or 48 h as indicated. When analysing stationary cells supplemented with calcium, cells were exposed to 100 mM calcium nitrate for 1 h after 24 h of growth.

For imaging, a Leica DMi8 inverted microscope was used, equipped with a Leica HC PL APO 100x/1,40 oil PH3 objective, CoolLED pE300W light source and a Leica K8 Scientific CMOS camera.Fluorescence of mNeonGreen and msfGFP was excited at using the CoolLED pE-300w light source in combination with a GFP filter cube (excitation 470/40 nm, dichroic 495 nm, emission 525/50 nm). For mNeonGreen, illumination intensity was set to 20% with an exposure time of 500 ms, while msfGFP was imaged at 2% intensity and 50 ms exposure time. Fluorescence of Nile Red was excited using the CoolLED pE-300w light source together with a TXR filter cube (excitation 560/40 nm, dichroic 585 nm, emission 630/75 nm) at 60% illumination intensity and 50 ms exposure time. Imaging was performed in biological triplicates. All images were analysed using ImageJ (Schindelin et al. 2012), while GFP intensity/µm2 was calculated from automatically detected cells using a custom Python script.

### Flow cytometry

Flowcytometry was used to quantify the GFP fluorescence intensity in reporter cell populations. The strain SGB1114 was analyzed in exponential and stationary growth phase. To reach the exponential phase, 5 ml of YA medium was inoculated in duplicate from an overnight culture in YA medium to a starting OD_600_ of 0.05 and incubated at 30°C with shaking (120 rpm). When the cultures reached an optical density of approximately 0.4, the culture was split into two 2.5 ml cultures and one batch was exposed to 100 mM calcium nitrate followed by incubation for another hour. The control culture was left untreated, with incubation continued the same as the challenged culture.

To prepare stationary phase cells, 5 ml of YA medium was inoculated in duplicate from an overnight culture to a starting OD_600_ of 0.05 and incubated at 30°C with shaking (120 rpm) for 24 h. Then, each culture was split into two 2.5 ml cultures and one batch was supplemented with 100 mM calcium nitrate followed by 1 h of additional incubation at 30°C. The other batch was further cultivated without the addition of calcium nitrate.

Prior to flow cytometry analysis, cultures were diluted at a 1:5 ratio with 0.85% (w/v) sodium chloride solution. The measurement was performed using the Attune NxT Flow cytometer (Invitrogen) equipped with 488 nm laser with a flow rate of 100 µl/min. Per sample, 50,000 single cells were measured. The GFP signal was detected in BL1 (530/30 nm). To set the threshold to detect GFP-positive cells above any intrinsic fluorescence of *S. silvestris* cells, a negative control strain was used which carried plasmid pMSMAD06, i.e. containing the gene encoding sfGFP but without a promoter.

### Bioinformatic analyses

The genomic information for *S. silvestris* was obtained from GenBank (accession number JBFEAN000000000), while the protein sequence and predicted structure of YloB from *B. subtilis* were retrieved from UniProt (accession number O34431). Reciprocal BLAST was performed using BLASTp algorithm (Altschul et al. 1990) against the *B. subtilis* 168 reference genome (RefSeq GCF_000009045.1). Protein and transmembrane domains were determine by HMMER scan (Potter et al. 2018) and 2D topology was analysed using PROTTER (Omasits et al. 2014) using default parameters. A pairwise alignment of YloB of *B. subtilis* and *S. silvestris* was conducted with ClustalW (Larkin et al. 2007) and conserved features were manually marked within the sequence based on the work of Raeymaekers, Wuytack et al. (2002). Protein structures were predicted with AlphaFold 3 using default settings (Abramson et al. 2024). Chimera X (Pettersen et al. 2021) was used to visualize structures and generate overlays between the predicted structures of *S. silvestris* and *B. subtilis* YloB proteins. encoding sfGFP but without a promoter.

## Supporting information

Numerical data

## Acknowledgements

We thank Elke Linden for technical support and for construction of pMSMAD06. We also thank Henrik Strahl and Jess Buttress, Newcastle University, UK, for expert advice and guidance on fluorescence microscopy and membrane staining.

