## Supplemental tables and figures for "Decoding the functional role of the calcium ATPase YloB in microbially induced calcite precipitation and sporulation in *Solibacillus silvestris*"

### Supplement

**Table S1.** Strains, vectors and plasmids used in this study

**Table S2.** Oligonucleotides used in this study

**Table S3.** Experimental numerical data available as separate Excel file.

**Figure S1.** The calcium ATPase from *S. silvestris* is a homolog of YloB from *B. subtilis*.

**Figure S2.** Structure comparison of YloB from *S. silvestris* and *B. subtilis*.

**Figure S3.** Representative growth curves with different calcium nitrate concentrations.

**Figure S4.** Cells lacking YloB show a sporulation defect after 24 h.

**Figure S5**. Gating strategy for flow cytometric analysis of $P_{\mathit{yloB}}$ *sfGFP*

**References**

**Table S1.** Strains, vectors and plasmids used in this study

| **Strains** | **Characteristic** | **Source** |
|---|---|---|
| E. coli ST18 | *RP4-2(Km::Tn7,Tc::Mu-1) pro-82* λpir *recA1 endA1 thiE1 hsdR17 creC510 ΔhemA* | (Thoma und Schobert 2009) |
| *S. silvestris* CGN12 | WT | (Reeksting et al. 2020) |
| SGB1106 | *S. silvestris*; Δ*yloB* | This study |
| SGB1108 | *S. silvestris; yloB::yloB-mNeonGreen* | This study |
| SGB1112 | *S. silvestris;* pMAD-oriT-*sfGFP* | This study |
| SGB1114 | *S. silvestris;* pMAD-oriT-$P_{ylo B}$ *sfGFP* | This study |
| SGB1117 | *S. silvestris;* Δ*yloB* pMAD-oriT-$P_{ylo B}$ *yloB* | This study |
| **Vectors** | | |
| pMAD-oriT | Temperature sensitive *ori*; $ery^R$; *oriT* for conjugation into *S. silvestris* | (Seidel et al. 2025) |
| pHJS105 | Contains genetic sequence for monomeric superfolder GFP | (Gamba et al. 2015) |
| pSHP1 | Contains genetic sequence for mNeonGreen | Henrik von Strahl, Newcastle University |
| **Plasmids** | | |
| pMSMAD03 | Δ*yloB* gene deletion; $ery^R$; upstream flank, truncated *yloB* and downstream flank | This study |
| pMSMAD05 | C-terminal mNG fusion to YloB; $ery^R$; upstream flank, *mNG* and downstream flank | This study |
| pMSMAD06 | Reporter assay control; $ery^R$; *sfGFP* without promoter | This study |
| pMSMAD08 | Reporter assay; $ery^R$; *sfGFP* under control of $P_{ylo B}$ promoter from *S. silvestris* | This study |
| pMSMAD11 | Complementation of Δ*yloB* mutant; $ery^R$; *yloB* gene under control of the native $P_{ylo B}$ promoter | This study |

**Table S2.** Oligonucleotides used in this study

| Primer | Sequence 5'-3'[a] |
|---|---|
| **pMAD-oriT cloning primers** | |
| SG1540 | gctaatgttacgttacacattaactagacag |
| SG1541 | gtctacacgaacccctttggc |
| SG1680 | gtcatatggatcccgcacgatatacaggattttgcc |
| SG1681 | gaattcgagctcccgggtaccatggc |
| **Construction of pMSMAD03** | |
| SG1682 | gccatttagcggcgtgtg |
| SG1683 | caaaaagcccggctgttcg |
| SG1684 | *gccttagtagacccttctgttactttc*atttgaaacctcctttaag |
| SG1685 | *gaaagtaacagaagggtctactaaggc*ataaacagaaaatag |
| SG1686 | gcatgccatggtacccgggagctcgaattccttttagtagcatcttcccacctctatg |
| SG1687 | caaaatcctgtatatcgtgcgggatccatatgacggatgacaatgctgagcaatatcg |
| **Construction of pMSMAD05** | |
| SG1694 | gcatgccatggtacccgggagctcgaattcgttgcagattgtattgaagcggg |
| SG1695 | *tgaaccaccaccacc*tgccttagtagacccgaaaaacag |
| SG1696 | *ggtggtggtggttca*atggtttcgaaaggagaggagg |
| SG1697 | *ttctattttctgtttacttatagagtt*catccatacccatcacg |
| SG1698 | *aactctataagtaaacagaaaatagaa*tgtataaaaatgcaaatggc |
| SG1699 | caaaatcctgtatatcgtgcgggatccatatgacgttgctttgtacttttctacgattgcttc |
| **Construction of pMSMAD06** | |
| SG1787 | *atcctgtatatcgtgcgggatccatatgac*ttatttgtagagctcatccatgccatg |
| SG1796 | gcatgccatggtacccgggagctcgaattcatgagcaaaggagaagaacttttcactg |
| **Construction of pMSMAD08** | |
| SG1790 | gcatgccatggtacccgggagctcgaattcctatcaaggctttgaaggtgtttttgc |
| SG1793 | atgagcaaaggagaagaacttttcactg |
| SG1795 | tccagtgaaaagttcttctcctttgctcatttgaaacctcctttaagtcatttgaatggtg |
| **Construction of pMSMAD11** | |
| SG1708 | gcatgccatggtacccgggagctcgaattcctatcaaggctttgaaggtgttttt |
| SG1820 | atcctgtatatcgtgcgggatccatatgacttatgccttagtagacccgaaaaacag |
| **Nested primers for *yloB* to screen for deletion/insertions** | |
| SG1842 | gccattgccgaactagaagaac |
| SG1843 | gaaactttgaaaccgtgtgctcc |

[a] overhangs for Gibson Assembly underlined, overhangs for overlap PCR in italics

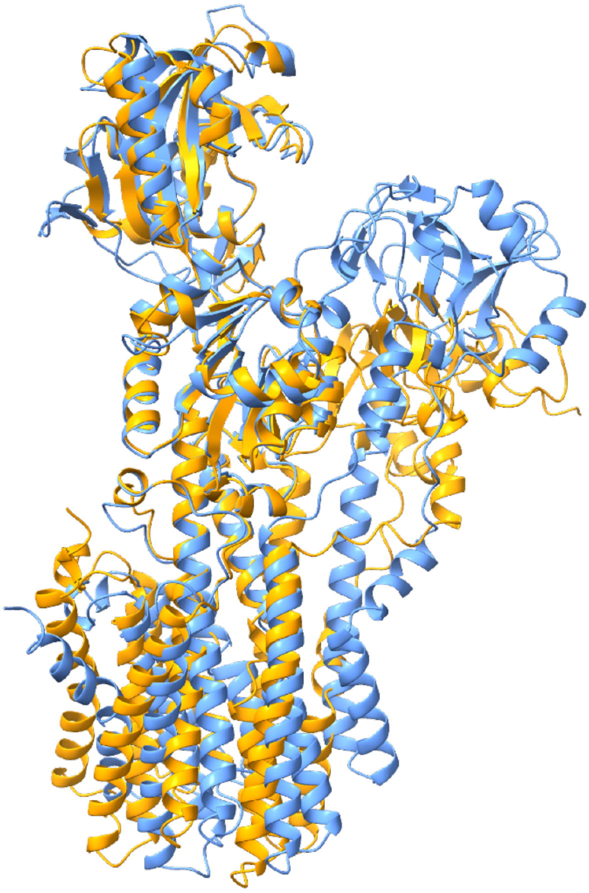

**Figure S1. Structure comparison of YloB from *S. silvestris* and *B. subtilis*.**
Superimposed models of the full-length YloB from *S. silvestris* (yellow) and *B. subtilis* (blue). Structure predictions were obtained from AlphaFold3 (Abramson et al. 2024) and visualized using ChimeraX (Pettersen et al. 2021).

```
B.subtilis     MKFHEMGQTDLLEATN--TSMKQGLTEKEVKKRLDKHGPNELQEGKKTSALLLFFAQFKD
S.silvestris   MKVTEYYQASVQEVMKKLDVTQYGLNDYEVRGRQKKYGYNELKEGKQKNIFQVFLEQFQD
               **. *  *:.: *. :     : **.: **: * .*:* ***:***:.. : :*: **:*

B.subtilis     FMVLVLLAATLISGFLGEYVDAVAIIAIVFVNGILGFFQERRAEQSLQALKELSTPHVMA
S.silvestris   FLVLILIVAAIVSIFLGDMDSSIVILVVIILNAILGTVQHARAEKSLNSLKEMSAPVSKV
               *:**:*:.*:::* ***:  .::.*:.::::*.***  *. ***:**::***:*:*   .
                     M1                    M2                 loop
B.subtilis     LREGSWTKIPSKELVPGDIVKFTSGDRIGADVRIVEARSLEIEESALTGESIPVVKHADK
S.silvestris   KRNGEILEIPSRNVIVGDLLILEAGDSISADGRVVESYSLQINESSLTGESLAVDKVANT
                *:*.  :***:::: **:: : :** *.** *:**: **:*:**:*****:.* * *:.

B.subtilis     LKKPDVSLGDITNMAFMGTIVTRGSGVGVVVGTGMNTAMGKIADMLESAGTLSTPLQRRL
S.silvestris   IHETELALGDRKNMVYSGSFVTNGRGAVAVTSIGMNTEIGKIAKLIDNAKEKKTPLQVNL
               :::.:::*** .**.: *::**.* *. .*.. **** :****.:::.*   .**** .*
                                                       ●  ● ●
B.subtilis     EQLGKILIVVALLLTVLVVAVGVIQGHDLYSMFLAGVSLAVAAIPEGLPAIVTVALSLGV
S.silvestris   DKFGKRLAIGIIFICIIIFALDIIRGREMIDSFMFAVSLAVAAIPEALSSIVTIVLAVGT
               :::** * :  ::: :::.*:.:*:*::: . *: .**********.*.:***:.*::*.
                            M3         phosph.        M4
                                          ▼
B.subtilis     QRMIKQKSIVRKLPAVETLGCASIICSDKTGTMTQNKMTVTHVWSGGKTWRVAGAGYEPK
S.silvestris   QNMAKENAIIRKLPAVESLGSVSIICTDKTGTLTQNKMTVQEIYTG-------------E
               *.* *:::*:*******:**..****:*****:******* .:::*             :

B.subtilis     GSFTLNEKEISVNEHKPLQQMLLFGALCNNSNIEKRDGEYVLDGDPTEGALLTAARKGGF
S.silvestris   QFFSSSELEPANPNHK---KLLDFAILCNDSMMLR----DKTIGDPTELALVKLGHEFEM
                 *: .* * :  :**   ::* *. ***:* : :        ***** **:. .::  :
                               ATP  ATP                FITC
                                ▼    ▼                  ▼
B.subtilis     SKEFVESNYRVIEEFPFDSARKMMTVIVENQDRKRYIITKGAPDVLMQRSSRIYYDGSAA
S.silvestris   SEQLIRGLHPRVGEIPFDSTRKLMSTVHRMGNRN-VMITKGAVDELLPRISRVDSS-SST
               *::::.. :  : *:****:**:*:.: .  :*:  :***** * *: * **:  . *::
                                                                       loop
B.subtilis     LFSNERKAETEAVLRHLASQALRTIAVAYRPIKAGETPSMEQAEKDLTMLGLSGIIDPPR
S.silvestris   IITNEQLEKINQANTAFSNAGLRVIAITYKDVFSSQIS--EKDEQGLTFVGLIAMMDPPR
               :::**:  : : .   ::. .**.**::*: : :.: .  *: *:.**::** .::****
                                 loop
B.subtilis     PEVRQAIKECREAGIKTVMITGDHVETAKAIAKDLRLLPKSGKIMDGKMLNELSQEELSH
S.silvestris   DESAQAVADCIEAGIKPVMITGDHKITAIAIAKQLGILKSPDEAIEGKEIEKLSDRELKN
                *  **: :* *****.*******  ** ****:* :* ...: ::** :::**:.**.:

B.subtilis     VVEDVYVFARVSPEHKLKIVKAYQENGHIVAMTGDGVNDAPAIKQADIGVSMGITGTDVA
S.silvestris   KVESFSVYARVSPEHKIRIVKAWQEKGHVVAMTGDGVNDAPALKQAEIGVAMGKTGTEVA
                **.. *:********::****:**:**:*************:***:***:** ***:**
                                                       ◆  ◆
B.subtilis     KEASSLVLVDDNFATIKSAIKEGRNIYENIRKFIRYLLASNVGEILVMLFAMLLALPLPL
S.silvestris   KDASAMILTDDNFLTIVKSISNGRSIYANIKNAIKFLLSGNAGAIFVVLYATLFALPAPF
               *:**:::*.**** ** .:*.:**.** **:: *::**:.*.* *:*:*:* *:*** *:
                                                         M5
                           ◆
                       ●  ◆●
B.subtilis     VPIQILWVNLVTDGLPAMALGMDQPEGDVMKRKPRHPKEGVFARKLGWKVVSRGFLIGVA
S.silvestris   LPIHLLFINLLTDSLPAIAIGLEPHNKKLMKEKPRHINESLLNKKFVSQVGFEGLIIAAV
               :**::*::**:**.***:*:*::  .  :**.****  :*.:: :*:  :*  .*::*...
                          M6                                           M7
                                              ◆
B.subtilis     TILAFIIVYHRNPENLAYAQTIAFATLVLAQLIHVFDCRSETSVFSRNPFQNLYLIGAVL
S.silvestris   TIIAFQVGLSTG--DTAVATTMAFATLCLSRLLHGFNCRSEESIFKIGIFSNLAVWIAFL
               **:** :    .  : * * *:***** *::*:* *:**** *:*. . *.** :  *.*
                                            M8
B.subtilis     SSILLMLVVIYYPPLQPIFHTVAITPGDWMLVIGMSAIPTFLLAGSLLTRKK---
S.silvestris   LGFTILNFVLING----LFEVANLTSNQYLMIYGLSLVPLVIIQVRKLFFGSTKA
                .: :: .*:       :*... :*..::::: *:* :* .::    *   .
                M9                              M10
```

**Figure S2. The calcium ATPase from *S. silvestris* is a homolog of YloB from *B. subtilis*.** The sequence alignment of the calcium ATPase from *S. silvestris* and *B. subtilis* was conducted using ClustalW. Features that are conserved and essential for functionality of P-type ATPases are annotated within the sequence. Residues of the first (orange, ●) and second (red, ◆) calcium binding sites, phosphorylation site (phosph., purple) and ATP binding sites (blue) are marked. Conserved loops, such as TGE loop located in the actuator domain, the MITGD loop, FITC-binding lysine and the DPPR loop connecting the nucleotide-binding and the phosphorylation domain, are highlighted (petrol). M1-10 are the predicted transmembrane helices. Conserved features were adapted from Raeymaekers et al. 2002. Identical amino acids are denoted by a star (*); a colon (:) represents conservation between amino acid groups of strongly similar properties; a period (.) indicates conservation between amino acid groups of weakly similar properties.

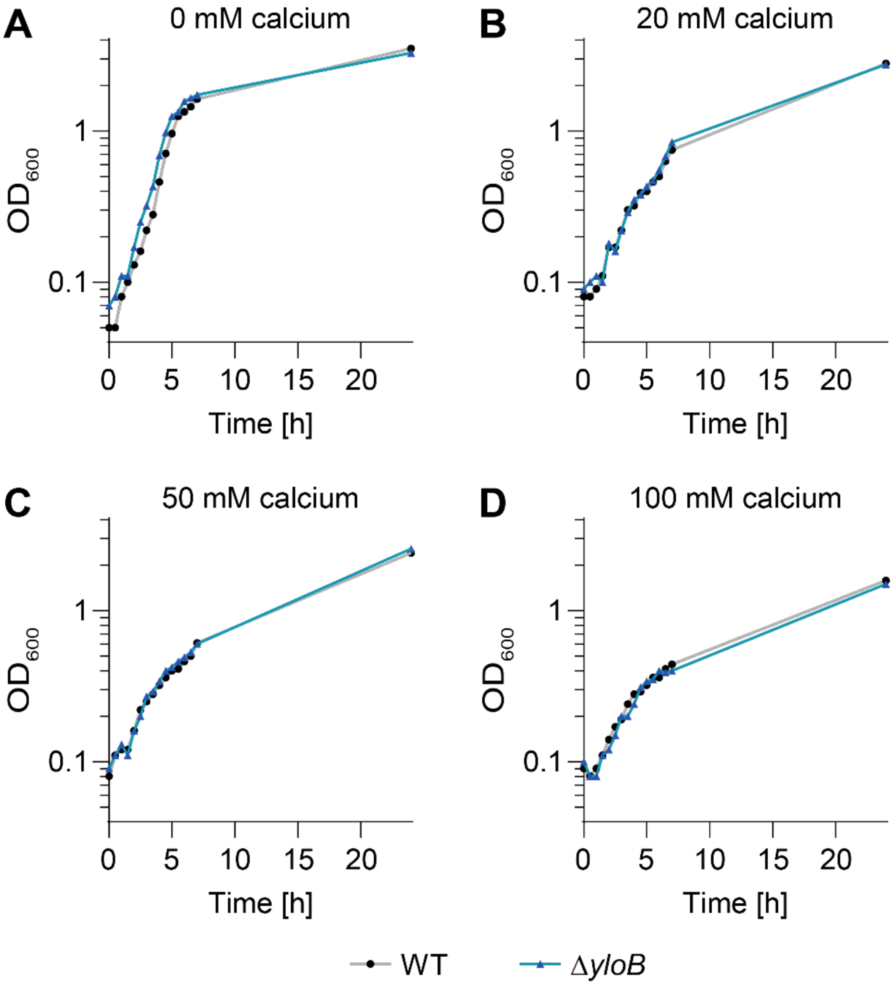


**Figure S3. Representative growth curves with different calcium nitrate concentrations.** YAC medium containing (A) 0 mM (B) 20 mM (C) 50 mM or (D) 100 mM was inoculated with either WT and Δ*yloB* and grown for 24 h, with $OD_{600}$ being measured every 30 min for the first seven hours and a final measurement after 24 h.

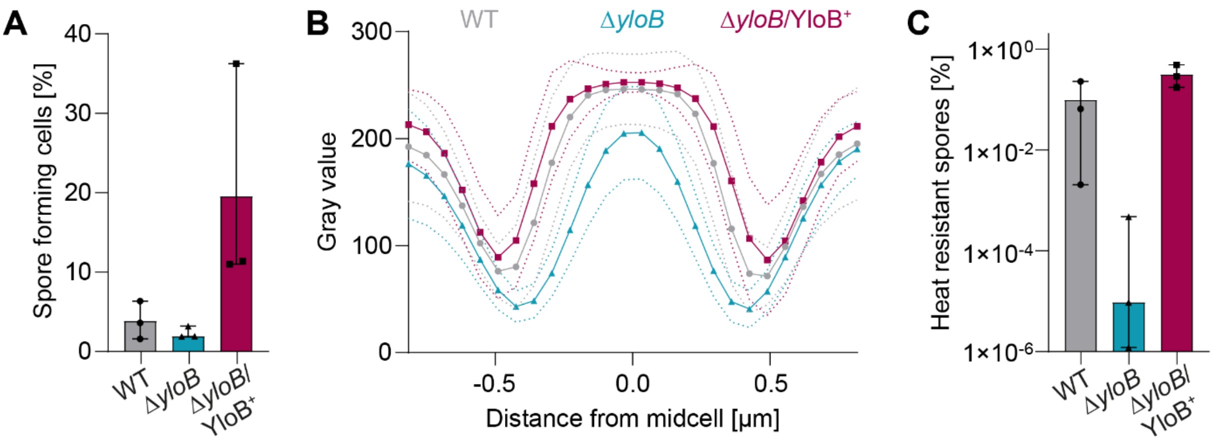


**Figure S4. Cells lacking YloB show a sporulation defect after 24 h.** (A) Proportion of total spores of indicated strains determined by microscopy. The fraction of sporulated cells was divided by the total cell count. The mean and the standard deviation for biological triplicates are represented. (B) Cells were grown in YA medium for 24 h. Microscopic images of sporulated cells for each strain are presented. Spore profile plot was determined representing the size and dehydration of the spore. The symbols depict the mean value, and the dotted line represents the standard deviation. (C) Ratio of viable spores was determined by dividing the CFU/ml of the pasteurized culture by the non-pasteurized sample. The mean and the standard deviation for biological triplicates are represented.

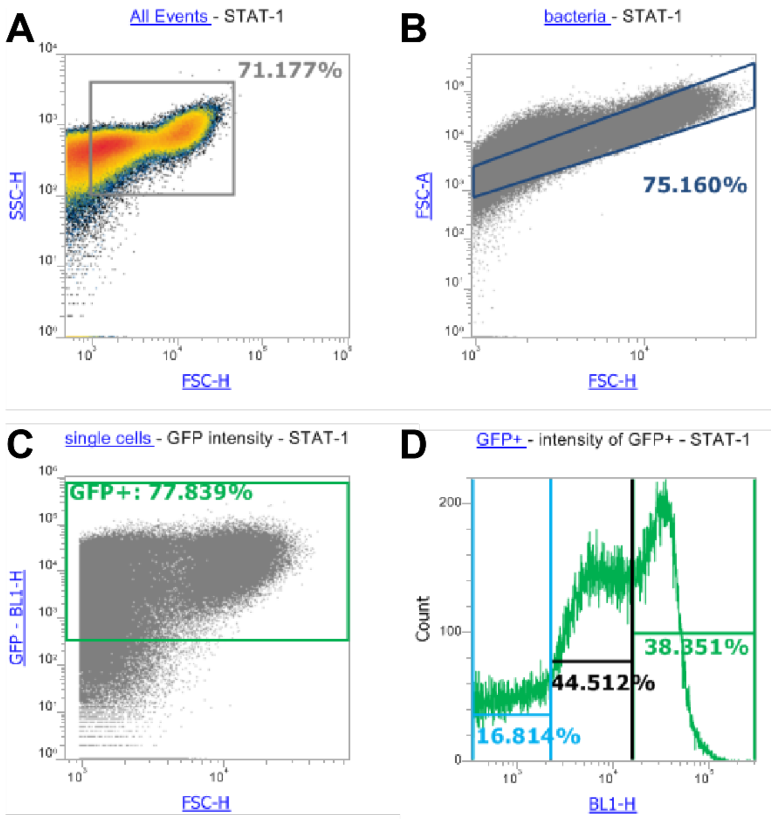


**Figure S5. Gating strategy for flow cytometric analysis of $P_{yloB}$ *sfGFP* activity.** Representative gating workflow for detecting single cells of strain SGB1114 producing msfGFP. (A) Bacterial cells were distinguished from debris based on forward scatter (FSC) and side scatter (SSC). (B) Single cells were identified using FSC-A versus FSC-H. (C) GFP signal was measured in channel BL1 and $GFP^+$ cells were identified using a threshold set using the promoter-less control strain SGB1112 as negative control. (D) $GFP^+$ cells were further subdivided into three fluorescence intensity levels (low, medium, high).
